# Crystal structure of chromo barrel domain of RBBP1

**DOI:** 10.1101/257527

**Authors:** Ming Lei, Yue Feng, Mengqi Zhou, Yuan Yang, Peter Loppnau, Yanjun Li, Yi Yang, Yanli Liu

## Abstract

RBBP1 is a retinoblastoma protein (pRb) binding protein acting as a repressor of gene transcription. RBBP1 is a multidomain protein including a chromo barrel domain, and its chromo barrel domain has been reported to recognize histone H4K20me3 weakly, and this binding is enhanced by the simultaneous binding of DNA. However, the molecular basis of this DNA-mediated histone binding by the chromo barrel domain of RBBP1 is unclear. Here we attempted to co-crystallize the chromo barrel domain of RBBP1 with either a histone H4K20me3 peptide alone or with both a histone H4K20me3 peptide and DNA, but only solved the peptide/DNA unbound crystal structure. Our structural analysis indicates that RBBP1 could interact with histone H4K20me3 similar to other histone binding chromo barrel domains, and the surface charge representation analysis of the chromo barrel domain of RBBP1 suggests that the chromo barrel domain of RBBP1 does not have a typical DNA binding surface, indicating that it might not bind to DNA. Consistently, our ITC assays also showed that DNA does not significantly enhance the histone binding ability of the chromo barrel domain of RBBP1.

## Introduction

RBBP1 (retinoblastoma-binding protein 1), also known as ARID4A, was first cloned from human complementary DNA expression library, which was found by using retinoblastoma protein (pRb, a tumor suppressor protein) as a bait to search for the potential interaction proteins of pRb [1]. As a corepressor of pRb, RBBP1 binds to the pocket domain of pRb and participates in the suppression of E2F (a transcription factor)-mediated transcription via both histone deacetylase (HDAC)-dependent and -independent repression activities [2,3]. RBBP1 functions as a tumor and leukemia suppressor and interacts with the mSin3A complex to repress gene transcription [4,5,6]. Further study also suggests that RBBP1 participates in regulation of epigenetic modification such as histone methylation as discovered in Prader-Willi/Angelman syndrome and leukemia [6,7].

RBBP1 is a multidomain protein, containing a double Tudor domain, a PWWP domain, an ARID domain, a chromo barrel domain and a R2 domain from the N terminus to the C terminus [8]. The ARID domain is reported to bind to DNA with AT-rich sequences, however the ARID domain of RBBP1 showed no DNA sequence preference [9]. Tudor domain, PWWP domain and chromo barrel domain are members of Royal family domains, which are reported to bind to the methylated histone tails [10,11,12,13]. However, the double Tudor domain of RBBP1, which forms an hybrid Tudor domain does not bind to methylated histone but interacts with DNA without sequential specificity [14]. The PWWP domain in general interacts with chromatin by synergistically binding to both histone and DNA [15]. However, the function of PWWP domain of RBBP1 is not reported yet. The chromo barrel domain of RBBP1 binds to histone H4K20me3 peptide at a millimolar affinity, and this binding is reported to be reinforced by the interaction of chromo barrel domain with DNA [8]. However, the molecular mechanisms of these interactions are still unclear.

To better understand the interactions between the chromo barrel domain of RBBP1 and histone H4K20me3 peptide and DNA, we aimed to analyze these interactions by quantitative binding assays and crystallographic analysis. In this study, we tried to analyze the interaction between the chromo barrel domain of RBBP1 and histone H4K20me3 peptide with or without DNA by quantitative isothermal titration calorimetry (ITC) binding assays, however the histone binding is too weak to be measured reliably and the addition of DNA does not enhance the histone binding. Next, we attempted to co-crystallize the chromo barrel domain of RBBP1 with both histone H4K20me3 peptide and DNA or with histone H4K20me3 peptide only, and solved the peptide/DNA unbound structure only. Further structural study indicates that chromo domain of RBBP1 could interact with H4K20me3 peptide similar to other chromo barrel domains, while it may not bind to DNA since it does not have a continuous positively charged surface like other DNA binding proteins.

## Results and discussion

### Interaction between chromo barrel domain of RBBP1 and histone H4K20me3 peptide not detected by ITC with or without DNA

According to a previous report, the chromo barrel domain of RBBP1 binds to H4K20me3 peptide and this interaction is strengthened by adding DNA [8]. Our isothermal calorimetry (ITC) assays failed to detect such binding to the H4K20me3 peptide with or without DNA (Fig. 1A and 1B). The reported binding affinity of the chromo barrel domain of RBBP1 to H4K20me3 is 6 mM [8], which might fall below our ITC assays’ limits of detection. However, we also could not observe higher binding affinity (0.4 mM) by adding DNA to the protein of the chromo barrel domain of RBBP1 before detecting its binding to histone H4K20me3 by ITC.

**Figure 1.**
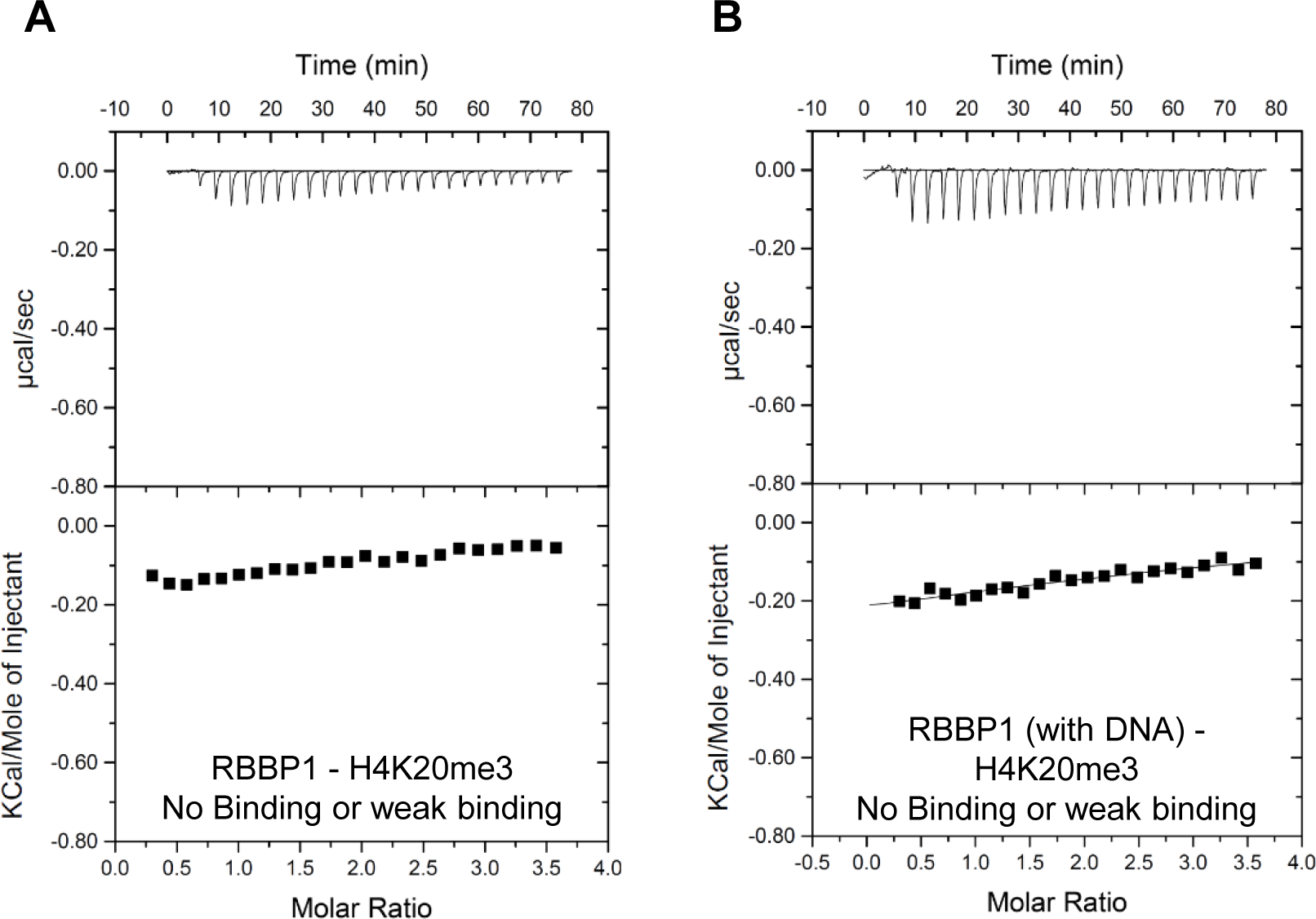
Interaction between chromo barrel domain of RBBP1 and histone H4K20me3 peptide not detected by ITC with or without DNA **(A and B)** ITC binding curves of histone H4K20me3 peptide to chromo barrel domain of RBBP1 without (A) or with DNA (B).

### Crystal structure of chromo barrel domain of RBBP1

To gain structural insight into the interaction between the chromo barrel domain of RBBP1 and its partners, we tried to crystalize it with its partners and solved the chromo barrel domain structure only (Table 1 and Fig. 2).

**Table 1.** Data collection and refinement statistics

|  |  |
| --- | --- |
| PDB entry | 6BPH |
| Data Processing |  |
| Space group | P43212 |
| Cell dimensions a, c [Å] | 41.5, 81.4 |
| Resolution (high resolution shell) [Å] | 36.97–1.80 (1.84-1.80) |
| Rmerge | 0.072 (1.077) |
| Mean(I/σ) | 26.2 (3.3) |
| Completeness [%] | 99.6 (98.8) |
| Multiplicity | 13.1 (13.3) |
| Model Refinement |  |
| Resolution [Å] | 36.90-1.85 |
| Reflections work / free | 6196 / 312 |
| R-value | 0.206 / 0.226 |
| Number of atoms: protein / water / unassigned (UNX) | 537 / 24 / 17 |
| Mean B-factor [Å<sup>2</sup>] protein / water | 30.8 / 37.7 |
| RMSD from ideal values: bonds [Å] / angles [°]<br>(PHENIX.MOLPROBITY) | 0.013 / 1.5 |

**Figure 2.**
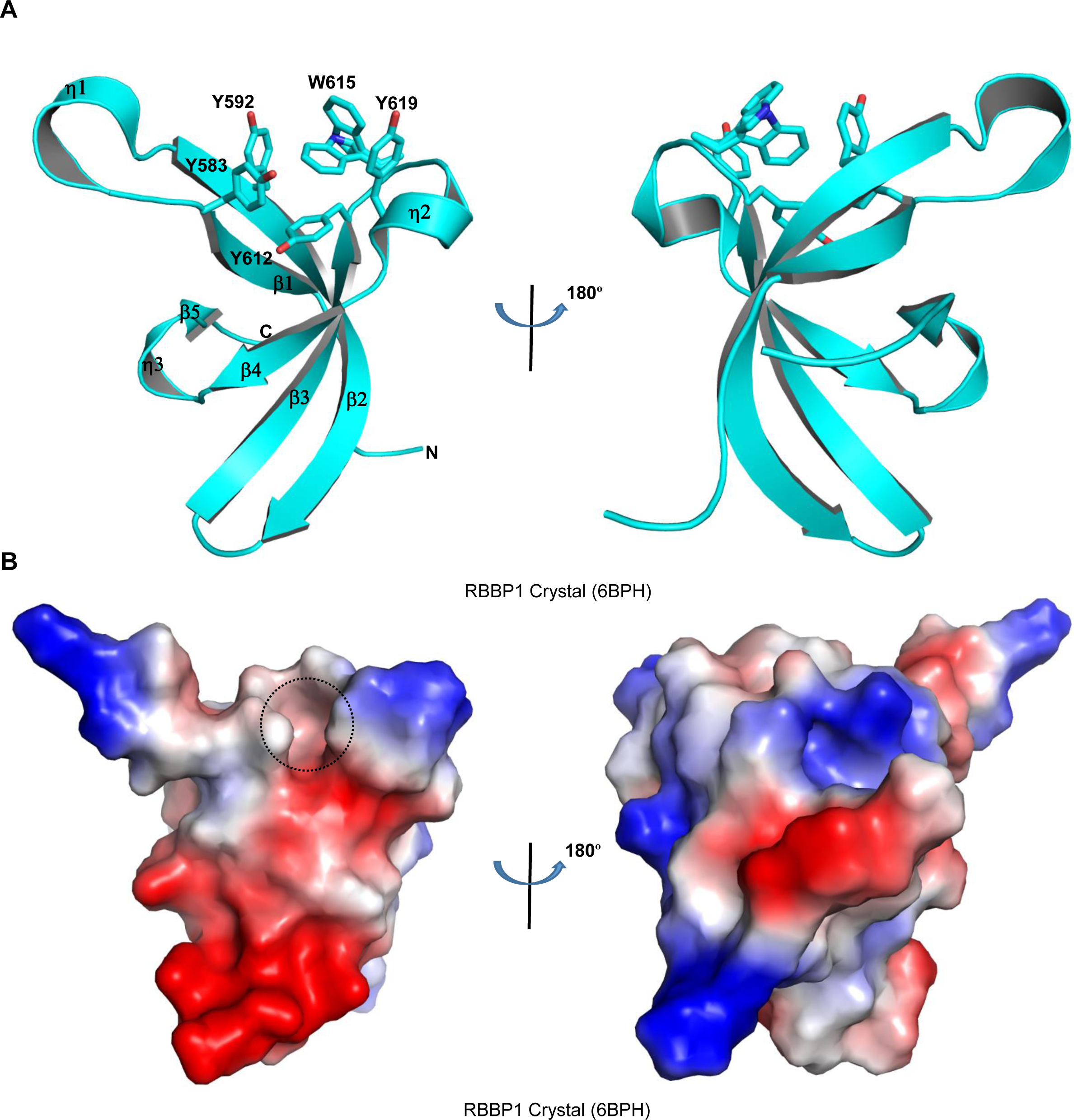
Structure of chromo barrel domain of RBBP1. **(A)** Overall structure of chromo barrel domain of RBBP1. The secondary structure elements are shown as cartoons and colored cyan, with the potential cage forming residues shown as sticks. **(B)** Electrostatic potential surface representation of chromo barrel domain of RBBP1 (isocontour values of ± 78.2 kT/e). Negative and positive potentials are depicted in red and blue, respectively. The potential binding cage resign for the methylated lysine is shown by black circle. Structure figures were generated by using PyMOL (http://pymol.sourceforge.net). Electrostatic potential surface representations were calculated with PyMOL’s built-in *protein contact potential* function [27].

There is one chromo barrel domain molecule in the asymmetrical unit in the RBBP1 crystal model. The overall structure of the RBBP1 chromo barrel domain is in accordance with the previously reported NMR structure (Fig. 3A) [8]. Briefly, it contains 5 β strands and 3 short 310 helices (η1, η2 and η3) with one between the β-strands of β1 and β2, one between the β-strands of β3 and β4, the other between the β-strands of β4 and β5 (Fig. 2A). The potential methylated lysine binding cage is made up by five aromatic residues, Tyr583, Tyr592, Tyr612, Trp615 and Tyr619, which side chains are oriented perpendicularly to each other to form five sides of a cube (Fig. 2A). The mutagenesis analyses have confirmed that residues Tyr592, Tyr612, Trp615 and Tyr619 play an important role in the H4K20me3 peptide recognition, however, Tyr583 is independent [8]. However, the electrostatic potential surface analysis indicates that there is no significant positively charged protein surface in the RBBP1 chromo barrel domain, which is consistent with its predicted theoretical pI value 5.76 (Fig. 2B). The protein surface of the RBBP1 chromo domain is different from other DNA binding proteins’ surface, such as the CXXC domains [16,17], therefore it may not bind to DNA.

**Figure 3.**
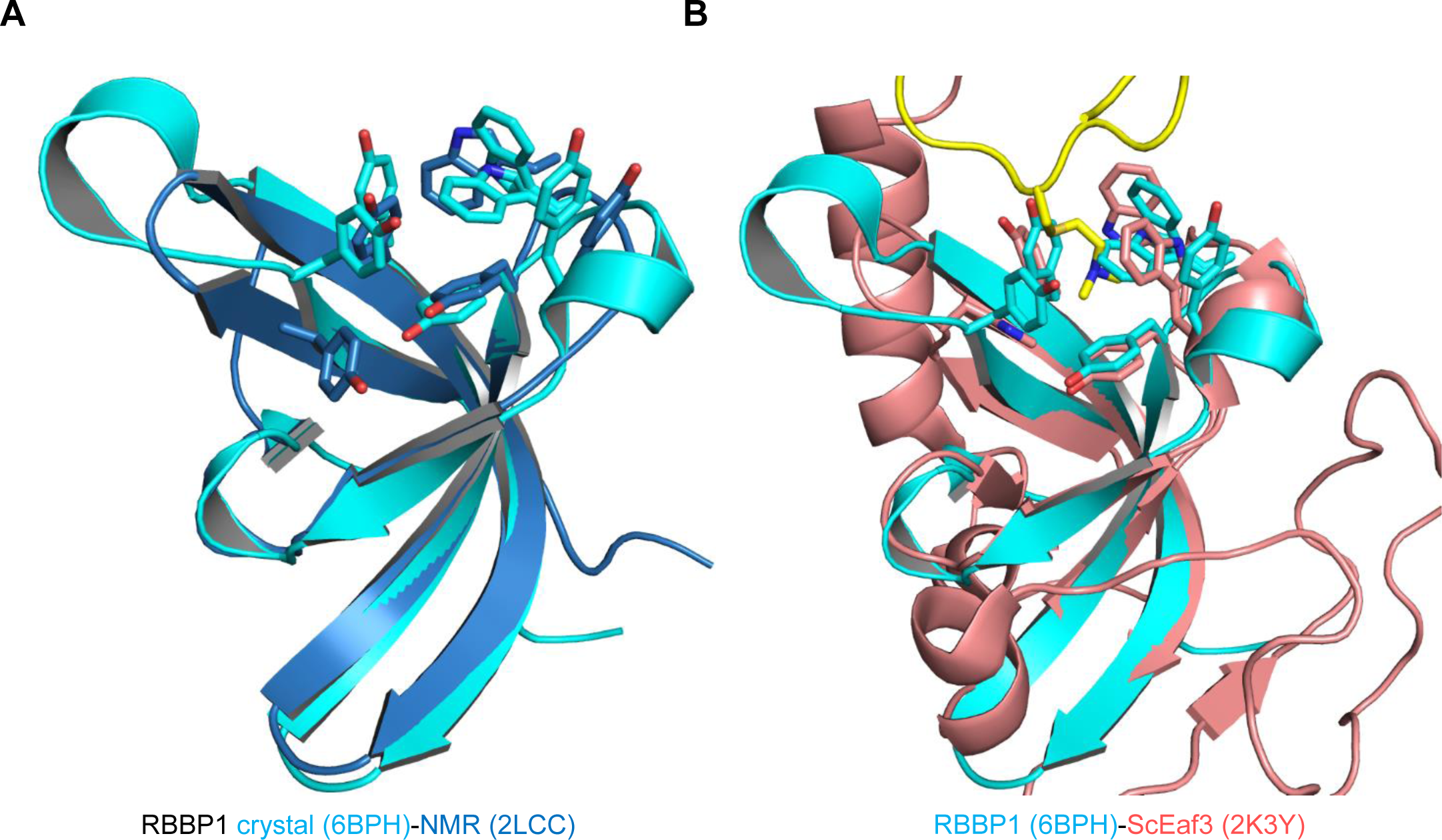
Structural comparison of chromo barrel domains. **(A)** Superimposition of chromo barrel domain of RBBP1 crystal structure (PDB code: 6BPH, cyan, this work) and NMR structure (PDB code: 2LCC, blue). **(B)** Superimposition of chromo barrel domain of RBBP1 (PDB code: 6BPH, cyan, this work) and ScEaf3 (PDB code: 2K3Y, salmon).

### Structure comparison of chromo barrel domain of TIP60 to other chromo barrel domain

The crystal and NMR structure of the chromo barrel domain of RBBP1 are almost the same, except two more short 310 helices (η1 and η2) in the crystal structure (Fig. 3A). Comparison to the ScEaf3 chromo barrel domain structure indicates that the C terminal long α helix is absent in the RBBP1 protein. The aromatic cage residues of RBBP1 are superimposed very well with those of the ScEaf3 protein, indicating that RBBP1 may recognize histone peptide in the same way as ScEaf3 (Fig. 3B).

In conclusion, our quantitative ITC binding assays reveal that the interaction between chromo barrel domain of RBBP1 and its partner is quite weak to be not reliably measured by regular ITC assay. Our crystal structure provides the molecular basis of the recognition of histone peptide by RBBP1 chromo barrel domain and indicates that it may not bind to DNA since it does not have a continuous positively charged surface like other DNA binding proteins, such as the CXXC domains.

## Materials and methods

### Protein expression and purification

The chromo barrel domain of RBBP1 (residues 568-635) was subcloned into a modified pET28-MHL vector. The encoded N-terminal His-tagged fusion protein was overexpressed in *Escherichia coli* BL21 (DE3) Codon plus RIL (Stratagene) cells at 15 °C and purified by affinity chromatography on Ni-nitrilotriacetate resin (Qiagen), followed by TEV protease treatment to remove the tag. Protein was further purified by Superdex75 gel-filtration (GE Healthcare, Piscataway, NJ). For crystallization experiments, purified protein was concentrated to 7.6 mg/mL in a buffer containing 20 mM Tris, pH 7.5, 150 mM NaCl and 1 mM DTT. The molecular weight of the purified protein was determined by mass spectrometry.

### Isothermal Titration Calorimetry (ITC)

The histone H4K20me3 peptide was synthesized by Peptide 2.0 Inc. For ITC measurements, the concentrated protein was diluted in 20 mM Tris, pH 7.5, and 150 mM NaCl. Likewise, the lyophilized peptide was dissolved in the same buffer and pH was adjusted by adding NaOH. Peptide concentration was estimated from the mass of lyophilized material. The two complementary ssDNA with sequences of 5’-ctcaggtcaaaggtcacg-3’, 3’-agtccagtttccagtgct-5’ (accoriding to previous study [8]) were dissolved in the same buffer and pH was adjusted by adding NaOH. Then the two ssDNA were annealed to dsDNA by PCR instrument by decreasing 5 °C/min from 95 °C to 25 °C. All ITC measurements were performed at 25 °C, using a VP-ITC microcalorimeter (GE Healthcare). Protein with a concentration of 100 μM or pre-mixed with DNA at a 1:1 molar ratio was placed in the cell chamber, and the peptide with a concentration of 2 mM in syringe was injected in 25 successive injections with a spacing of 180 s and a reference power of 13 μcal/s. Control experiments were performed under identical conditions to determine the heat signals that arise from injection of the peptides into the buffer. Data were fitted using the single-site binding model within the Origin software package (MicroCal, Inc.).

### Crystallization

Purified RBBP1 (7.6 mg/mL) was mixed with dsDNA and H4K20me3 peptide at 1:1.2:5 molar ratio, and crystallized using the sitting drop vapor diffusion method at 18 °C by mixing 0.5 μL of the protein with 0.5 μL of the reservoir solution. The crystals were obtained in a buffer containing 25% PEG 3350, 0.2 M magnesium chloride, 0.1 M Hepes, pH 7.5.

### Data collection and structure determination

Diffraction data were collected on a copper rotating anode x-ray source under sample cooling to 100 K. Diffraction images were processed with XDS [18] and symmetry-related intensities merged with AIMLESS [19]. The structure was solved by molecular replacement with PHASER [20] software and coordinates from PDB entry 2LCC [8]. The model was automatically rebuilt with ARP/wARP [21] and interactively modified in COOT [22]. REFMAC [23] and AUTOBUSTER [24] were used for restrained model refinement. PHENIX [25] programs were used for geometry validation [26] and calculation of model statistics. Additional information on crystallographic experiments and models is listed in Table 1.

### Accession number

Coordinates and structure factor amplitudes of chromo barrel domains of RBBP1 were deposited in the PDB with accession code 6BPH.

## Acknowledgements

We would like to thank Wolfram Tempel for data collection and structure determination. Results shown in this report are derived from work performed at Argonne National Laboratory, Structural Biology Center at the Advanced Photon Source. Argonne is operated by UChicago Argonne, LLC, for the U.S. Department of Energy, Office of Biological and Environmental Research under contract DE-AC02-06CH11357. This study was supported by the National Natural Science Foundation of China [grant number 31500613] and Hubei Chenguang Talented Youth Development Foundation. The SGC is a registered charity (number 1097737) that receives funds from AbbVie, Bayer Pharma AG, Boehringer Ingelheim, Canada Foundation for Innovation, Eshelman Institute for Innovation, Genome Canada through Ontario Genomics Institute [OGI-055], Innovative Medicines Initiative (EU/EFPIA) [ULTRA-DD grant no. 115766], Janssen, Merck KGaA, Darmstadt, Germany, MSD, Novartis Pharma AG, Ontario Ministry of Research, Innovation and Science (MRIS), Pfizer, São Paulo Research Foundation-FAPESP, Takeda, and Wellcome.

